# Viewing geometry and pattern type shape the detectability of snakes

**DOI:** 10.64898/2026.10.12.757824

**Authors:** Zixiang Li, Laura A. Kelley, Martin Stevens

## Abstract

Protective coloration can reduce predation risk by altering how prey are detected, recognised, or attacked. High-contrast markings may function as warning signals, camouflage, or both. In snakes, dorsal colour patterns are highly variable but the function remains debated, particularly because their appearance to an observer may change with viewing distance and angle. This study tested whether the function and efficacy of common snake pattern types (blotches, spots, diamonds, zig-zags and bands) is observer-view dependent, mediated by viewing distance and angle. Artificial snake models marked with these different pattern types that are broadly common across snake taxa, were placed in woodland habitat with a mixed shrub and grass understorey in Cornwall, UK, with photographs taken for evaluating levels of predicted camouflage and contrast against the background. Next, human volunteers searched for models, and we recorded detection distances and angles. Field results showed no significant difference in overall detection rates between pattern types; however, zigzag-patterned models were more likely to be detected at significantly shorter distances than blotch-patterned models, with a marginal trend towards shorter distances than diamond-patterned models, indicating effective longer-range camouflage. Detection risk did not alter gradually as volunteers approached but increased abruptly, correlating with image analyses showing that levels of predicted disruptive camouflage decreased with reduced distance. When detection was analysed using viewing angle, an advantage of zigzag patterns was robust to geometric scale, and observer height may further shape detection through viewing geometry. These findings suggest that snake dorsal patterns could potentially serve dual roles in reducing detection from afar, while being more conspicuous at close range. In addition, the results highlight the need to consider three-dimensional viewing geometry, including distance and visual angle, when evaluating the ecological function of animal coloration.

## Introduction

Animals evolve different kinds of protective coloration to influence both detection and perception by predators (Cott, 1940; B. Endler, 1991). Crypsis and aposematism are two common forms of protective coloration, traditionally thought to occupy opposite ends of an evolutionary spectrum of prey conspicuousness, with a cryptic target being inconspicuous and an aposematic target expected to be highly visible. Crypsis and aposematism are not necessarily mutually exclusive, however, but rather form a continuum from cryptic to conspicuous (Stevens, 2007), and recent work indicates that the same patterns may even play a role in multiple strategies (Barnett, Michalis, Scott-Samuel, & Cuthill, 2018). Some patterns can serve dual functions, allowing prey to navigate the trade-offs between background matching and predator avoidance. This is especially relevant in patchy habitats, where closely matching all microhabitats is difficult (Merilaita, Tuomi, & Jormalainen, 1999). For example, camouflage and aposematic signalling can be combined through distance-dependent coloration, whereby the same pattern is relatively cryptic at longer distances but more conspicuous at close range. Research using artificial prey shows that combining these functions can enable prey achieve higher survival than purely cryptic or conspicuous forms (Barnett & Cuthill, 2014; Barnett et al., 2016).

Animals often use high-contrast markings as warning signals (Stevens & Ruxton, 2012). However, such patterns may also contribute to camouflage through disruptive coloration and distance-dependent background matching, enabling crypsis even when animals with contrasting pattern elements do not closely match the background. For example, aposematic marginal wing markings in the wood tiger moth can provide disruptive camouflage (Honma, Mappes, & Valkonen, 2015). Due to the limitations of animal visual acuity, camouflage can be achieved through patterns with spatial frequencies that cannot be visually resolved from a distance (Barnett & Cuthill, 2014), by optically blending their pattern and body colour into the average background colour (Barnett, Cuthill, & Scott-Samuel, 2017; Barnett et al., 2016). This can enable background matching at a distance, but conspicuousness when up close (Guilford & Dawkins, 1991; Tullberg, Merilaita, & Wiklund, 2005). Conversely, disruptive coloration breaks up the body’s edges and contours through relatively high contrast internal and marginal patterns, creating false edges that can interfere with a predator’s ability to detect or recognize prey based on shape or outline cues (Cott, 1940; Stevens & Merilaita, 2009; Thayer, 1909).

For an irregular three-dimensional target, the perceived image should change with the observer’s position, meaning that the effectiveness of a camouflage strategy may vary significantly as viewing angle shifts during a predator’s approach (Hancock et al., 2023; Li, Hancock, Kelley, & Stevens, 2026). However, current studies are largely confined to specific viewing angles (Barnett, Cuthill, & Scott-Samuel, 2018; Bohlin, Tullberg, & Merilaita, 2008; Siddiqi, Cronin, Loew, Vorobyev, & Summers, 2004; Tullberg et al., 2005). In natural environments, changes in observer height can alter the relative position of the target against the background and affect the degree of obstruction by vegetation, potentially affecting recognition accuracy (Hancock et al., 2023). Baker et al. (2024) also found that viewing angle affected humans’ ability to distinguish Batesian mimics in hoverflies, although the effect was species-specific. Because observer distance changes are usually accompanied with viewing angle changes, whether viewing angle affects the effectiveness of certain camouflage patterns and whether camouflage effectiveness is angle-dependent remains to be explored.

Snakes exhibit a wide variety of patterns, which may serve various functions, from camouflage to warning signals (Allen, Baddeley, Scott-Samuel, & Cuthill, 2013; Jackson, Ingram, & Campbell, 1976; Li et al., 2026). Previous studies indicate that these patterns may serve a camouflage function through background matching or disruptive coloration (Creer, 2005; Isaac & Gregory, 2013; Wilson, Heinsohn, & Wood, 2006), while others show support for a possible aposematic function (Brodie III & Janzen, 1995; J. Valkonen, Niskanen, Bjorklund, & Mappes, 2011; J. K. Valkonen, Vakkila, Pesari, Tuominen, & Mappes, 2020). Some high-contrast snake patterns may therefore combine camouflage and warning functions (Li et al., 2026; J. K. Valkonen et al., 2020), but how these effects vary with distance and viewing angle remains unclear. Additionally, snakes face predation threats from a broad range of viewing elevations, from low terrestrial viewpoints to more elevated terrestrial and aerial viewpoints (Speybroeck, Beukema, Bok, & Van Der Voort, 2016). This makes the snake patterns particularly suited to test angle-dependent detectability, as patterns that disrupt an outline laterally may do so less effectively from steeper angles.

Although image-based analyses suggest that snake markings can generate context-dependent visual effects across habitats, viewing distances and viewing angles, whether these effects translate into detectability during active search remains unclear (Li et al., 2026). Here, we tested this by conducting a field experiment in which human observers searched for artificial snake models, with detection distance and viewing angle recorded, alongside analysing images of these models to quantify estimated camouflage levels. We predict that (1) detectability would differ among the five snake pattern treatments, reflecting differences in their potential camouflage functions; (2) models would generally be less detectable at longer distances, and become increasingly likely to be detected as observers approached; and (3) detectability would vary with viewing geometry, specifically viewing distance and viewing angle.

## Methods

### Artificial Model Preparation

We generally followed the methods for model creation of Li et al. (2026). The artificial models were not intended to mimic any particular snake species, but instead represented broadly occurring snake colour and pattern types. Replicas of snakes approximately 25 cm in length and 2.5 cm in diameter, were crafted using soft rubber snake models (Cobee). Thirty rubber snakes (three models for each colour-pattern group) were painted using spray paint in brown (151 Products Ltd, brown) and grey (Paint Factory Ltd, light grey). Five distinct pattern types were hand-painted in black on these replicas (Figure 1a), and their shapes and sizes approximately estimated according to photos of real snakes: (a) blotches, irregular patches spread haphazardly on the model; (b) spots, two rows of paired spots on the lateral side of the body (diameter 1.0±0.2 cm; spacing approximately 0.1 cm); (c) diamonds, a chain of round spots on the dorsal surface (diameter 1.0±0.1 cm; spacing approximately 0.1 cm) ; (d) zigzags, dorsal zigzag design with successive elements spaced approximately 2 cm apart; and (e) banding, circular stripes spaced evenly along the length of the model with 1.0±0.1 cm wide and separated by intervals of approximately the same width.

**Figure 1.**
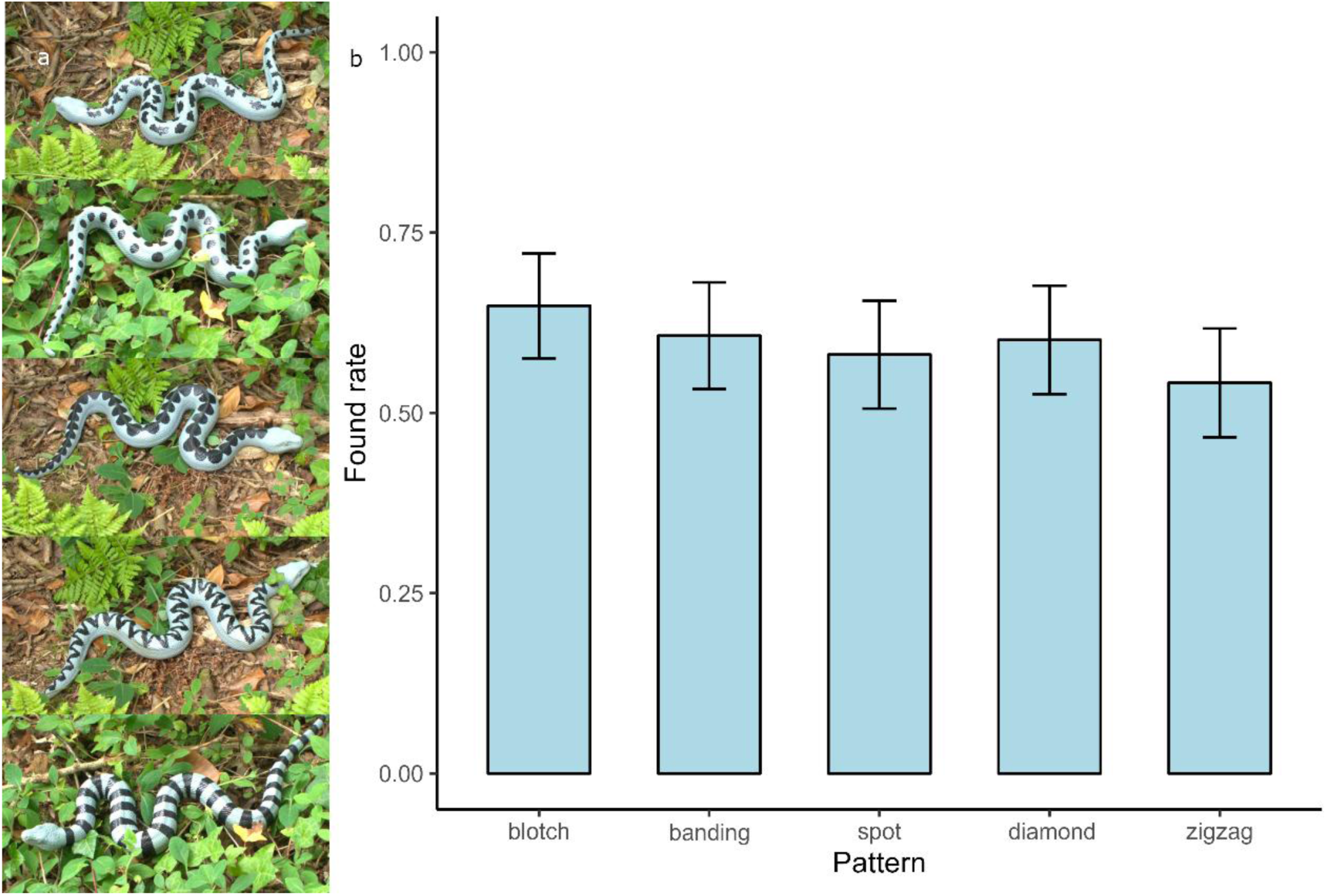
(a) Five pattern kinds on grey snake replicas, from top to bottom are blotch, spot, diamond, zigzag, and banding pattern. (b) Detection rates for different snake patterns (blotch: 0.648 ± 0.037, N=165; banding: 0.607 ± 0.038, N=168; spot: 0.581 ± 0.038, N=167; diamond: 0.601 ± 0.038, N=163; zigzag: 0.542 ± 0.038, N=168). Bar plots show the mean detection rate with Wald 95% confidence intervals based on binary survival status (0/1) observations.

The five pattern types were selected to represent broadly occurring forms of snake patterning rather than the coloration of particular species. Their shapes and relative dimensions were approximately based on patterns observed in photographs of real snakes. The blotch pattern served as the baseline because it is a common snake pattern and a reference category in comparative work (Allen et al., 2013). The spot pattern and diamond pattern represented different distributions of discrete markings, and the zigzag pattern and the banding pattern represented the different arrangements of transverse markings. The pattern treatments were whole-pattern phenotypes rather than isolated manipulations of a single visual feature. We standardised the approximate size and longitudinal spacing of repeated markings along the dorsal midline, but shape, orientation, black coverage and apparent coverage necessarily varied among treatments and with viewing angle. These differences, including associated brightness variation, were treated as components of each pattern phenotype.

### Research Sites and model placement

The study was conducted from 20th June to 18th September 2024 at College Reservoir (50°9’37 N, 5°7’22 W) in Penryn, Cornwall, UK. The reserve contains woodland with a dense shrub and grass understorey, adjacent to wetland and meadow habitats. At the regional level, Cornwall supports two native snake species: the adder, *Vipera berus*, and the barred grass snake, *Natrix helvetica*. A transect was set along a footpath, with 30 placement points randomly selected alongside the transect within two meters, on the left or right hand side of the transect line in equal numbers. The snake models were repositioned between each participant to randomize the sequence. At each point, the snake models were selected haphazardly by blindly choosing a model from a bag and directly placing it on the ground. As in Briolat, Arenas, Hughes, Liggins, and Stevens (2021), each point was at least 10 meters apart to prevent volunteers from searching for more than one model at once. The models were also ensured to be visible from this distance without full vegetation obstruction.

### Photography and volunteer search

At the beginning of the experiment, photographs were taken at each of the 30 placement points for all five pattern types in brown and grey colour variants (300 photographs in total). A digital camera was positioned approximately 1.5 meters and at a 45 degree angle from the subject under natural light. Each photograph included an 18% grey standard card and a ruler for image and scale calibration (Stevens, Párraga, Cuthill, Partridge, & Troscianko, 2007; Troscianko & Stevens, 2015) for calibration of the images. Images were captured in RAW format using a Sony A7 digital camera with a 28–70 mm lens, at approximately 1.5 m from the model and an angle of approximately 45° under natural light. Aperture was set to f/6.3 and ISO to 160.

From 23 Aug to 19 Sept in 2024, 28 volunteers (aged 18 to 35, 14 males, 14 females) with normal or corrected to normal vision and no known colour blindness participated. Volunteers were recruited through oral conversation and email. Each volunteer undertook the search individually. Before starting, a distance measure (Zamo 4th gen, Bosch, Germany) was given to each volunteer for measuring distance to the model. Participants’ prior familiarity with British snakes, the local snake fauna or the study site was not recorded, and they therefore could not be classified as either snake-familiar or snake-naïve. However, all participants were shown the same reference images of the experimental treatments before searching, thereby standardising their immediate familiarity with the target appearances. The height of volunteers ranged from 155 to 188 cm, and was recorded as an approximation of eye height to nearest cm for calculating the visual angle.

The researcher walked 3 m ahead of the volunteer to control pace without influencing volunteer search. Volunteers were encouraged to keep the same and comfortable speed rather than stopping to scan. When a target was spotted, the volunteer stopped and identified the pattern type, then measured the distance to the model. For analysis, the two detections that slightly exceeded the 10-m starting distance (10.3 m and 10.4 m, both from one volunteer) were truncated to 10 m to match the upper limit of the approach scale used in the survival analysis. The distance meter was held approximately at upper-abdomen height. For the geometric calculation of viewing angle, the height of the distance meter was approximated as 60% of the observer’s standing height, and 0.4 m was added to the measured distance as an approximate correction for forward arm extension. If a snake model was missing from its original spot (e.g. moved by people or animals), it was recorded as missing and not included in the statistics. The weather conditions were also recorded for each detection event and classified as overcast or sunny, depending on whether the objects could form distinct shadows.

### Image analysis

All image processing and analyses were performed using the Multispectral Image Calibration and Analysis Toolbox v2.2.2 (Troscianko & Stevens, 2015) in ImageJ 1.53v (Schneider, Rasband, & Eliceiri, 2012). Images were calibrated using a grey standard to normalize lighting conditions and saved as 32-bit multispectral images. Subsequently, scale bars were added based on the scale present in each photograph. The snake model outline in each image was manually selected as the region of interest (ROI), excluding background obstacles unless they completely obscured the snake’s outline. A bounding circle encompassing the ROI defined the local background, while the other part of ground background determined the global background for habitat metric calculations. Following Li et al. (2026), acuity-corrected (Caves & Johnsen, 2018) views with distance scaling were applied in the image analysis to account for the distance-dependent blur, using human visual acuity as 72 cycles per degree (cpd) (Land & Nilsson, 2012). Distance modelling ranged from 5 to 15 meters in 2.5 meter increments.

Colour congruence between the target snakes and their respective backgrounds using a logarithmic form of the receptor noise model (RNL) (Vorobyev & Osorio, 1998), with contrasts expressed in just noticeable difference (JND). ΔS is the discrimination threshold which is determined by the quantum catch and the respective noise level of the receptor channels. Generally, a JND value less than 1 is considered indistinguishable under ideal viewing conditions, while increasing values above this suggest increasingly discernible differences (Perez-Martinez, Riley, & Whiting, 2020; Siddiqi et al., 2004; Stevens, Troscianko, Wilson-Aggarwal, & Spottiswoode, 2017). For human colour vision, we used a Weber fraction of 0.05 with cone ratio X:Y:Z = 0.629:0.314:0.057 (Hofer, Carroll, Neitz, Neitz, & Williams, 2005). Achromatic target–background contrast (ΔS_Lum) was calculated from the luminance channel, which was defined as the mean of the LWS and MWS cone catches, in JNDs using a Weber fraction of 0.11 (Hofer et al., 2005; Troscianko & Stevens, 2015). Because treatments differed in black and apparent coverage, ΔS_Lum was used to assess whether luminance contrast might contribute to detectability. As luminance was not independently manipulated, it was interpreted descriptively rather than as an isolated causal effect.

Potential edge disruption was quantified in the luminance channel using Gabor filtering, which measures false edges orthogonal to the true outline (Troscianko et al., 2017). We used a luminance Weber fraction of 0.11, GabRat σ = 3, the human visual system, and the same distance scaling described above. Higher GabRat values indicate greater pattern–outline interference; values below 0.2 suggest weak disruption (Troscianko et al., 2017, Price et al., 2019). The image analyses quantify appearance under standardized photographic conditions, rather than reconstructing the exact visual scene at each field detection event. Because high GabRat can also arise from high-contrast (e.g. aposematic) patterns, its interpretation as evidence for camouflage function relies on the performance in the volunteer detection test.

### Statistical Analysis

All statistical tests were accomplished in R v4.2.3 (R Core Team, 2021). Raw detection frequencies among pattern types were first compared using a Pearson chi-square test. Then, for testing if pattern affects detection, the detection rate between treatment types was compared with a generalized linear mixed-effect model (GLMM) with a binomial distribution and logit link function, with pattern (blotches, spots, diamonds, zig-zags, bands), colour (brown, grey), and weather (sunny, overcast) as fixed effects. Random effects included point location, volunteer ID, and search duration. The inclusion of duration as a random effect accounts for between-trial variation in walking speed and search pace that could otherwise confound detection probability.

To analyse how detection risk changed as observers approached the target, we used piece-wise exponential additive mixed models (PAMMs). Although PAMMs are a time-to-event framework, here distance progressed from the starting point towards the target was used in place of actual time. Detection hazard therefore represents the rate of first detection at a given point in the approach. Detection status was used as the event variable, detection distance was converted to an approach scale by subtracting it from 10 m, so that increasing values represented movement towards the target; models that remained undetected were right-censored at the end of this scale. The data were then converted into piece-wise exponential data using the package pammtools (Bender et al., 2018). The continuous approach scale was divided into intervals, with interval boundaries based on quantiles of observed event times. Models were fitted using the package mgcv (Wood, 2011) with Poisson error distribution. The primary distance-based PAMM included pattern, colour, and weather as fixed effects, with random intercepts for volunteer and point using smooth terms with random-effect bases. A smooth term of the approach scale described the non-linear changes in detection hazard shared across snake patterns. We compared this with an extended model containing pattern-specific smooth terms for the approach scale within each pattern treatment, which allowed the shape of the hazard trajectory to differ among patterns. Pairwise contrasts among pattern treatments were estimated using emmeans (Lenth, 2023). As a sensitivity analysis, we ran the same model using angle as an alternative time scale to assess the robustness of the pattern effect to viewing geometry. To examine whether observer height contributed to detection through viewing geometry, supplementary distance-based models included a smooth effect of height and subsequently a tensor-product interaction between approach scale and height to test whether their effects were non-additive. Models were compared using AIC.

For the image analysis, linear mixed models (LMM) fitted with restricted maximum likelihood were chosen to analyse GabRat value, ΔS and ΔS_Lum separately, with pattern, colour and their interactions treated as fixed effects, point ID was included as a random effect, mitigating the influence of point selection. ANOVA was used to determine the significance of effects, followed by post-hoc tests (Lenth, 2023). The differences between adjacent distance levels within the same pattern group were measured by t-tests. The result of the picture analysis was compared with the volunteer search result to check consistency between estimated camouflage levels and the field experiment.

### Ethical Note

This study involved adult human participants searching for artificial snake models; no live animals were used. The complete protocol, including volunteer recruitment and testing, the participant information sheet and the written consent form, was reviewed and approved by the authors’ faculty-level institutional research ethics committee (Application ID 528352; committee name and institutional identity withheld for double-anonymous review). Before participation, volunteers received written information about the study procedures and provided written informed consent to participate and for their anonymized data to be analysed for research purposes. Participation was voluntary, and participants could withdraw without penalty. The task involved no deception or potentially harmful stimuli and presented no more than the minimal risks associated with walking in an outdoor public reserve. Participant data were anonymized before analysis, and no personally identifying information is reported. The research was conducted in accordance with the Declaration of Helsinki and its later amendments. Permission to conduct the fieldwork was granted by the organisation responsible for managing the study site.

## Results

### Detection rate

Of 840 targets presented across 28 trials, nine were lost and excluded; in the remaining 831 cases, 495 (59.57%) were detected and 336 (40.43%) were not detected. The average detection distance for volunteers was 2.42 m, N=495. Overall detection rate did not differ among patterns (Pearson’s chi-square test: χ²_4_ = 4.211, P = 0.378, N=831; Figure 1b), although the zigzag coefficient relative to blotch was negative but marginally non-significant. grey models were more detectable, whereas direct sunlight reduced search success (Table 1).

**Table 1.** Effects of fixed effects on detection success in GLMM model.

|  | Estimate Std. | 95% CI | z value | Pr(>\|z\|) |  |
| --- | --- | --- | --- | --- | --- |
| (Intercept) | 1.269 | [0.667, 1.871] | 4.136 | **<0.001** | *** |
| Pattern: Banding | -0.258 | [-0.773, 0.257] | -0.980 | 0.327 |  |
| Pattern: Spot | -0.321 | [-0.829, 0.187] | -1.240 | 0.215 |  |
| Pattern: Diamond | -0.268 | [-0.778, 0.242] | -1.033 | 0.302 |  |
| Pattern: Zigzag | -0.508 | [-1.018, 0.002] | -1.950 | **0.051** | . |
| Colour: Grey | 0.601 | [0.197, 1.005] | 2.914 | **0.004** | ** |
| Weather: Sunny | -1.228 | [-1.700, -0.756] | -5.091 | **<0.001** | *** |

### Detection risk through distance change

During the volunteer search process, detection risk changed non-linearly as observers approached the target. In the distance-based PAMM, the baseline smooth term was highly significant (Table 2), indicating that detection hazard did not increase linearly with the approach process. Instead, the probability of detection changed sharply over part of the search trajectory, consistent with a threshold-like pattern in detectability. Volunteer identity and model placement point also explained variation in detection risk, as indicated by the significant random-effect smooths for volunteer and point.

**Table 2.** Model summaries for distance-based PAMMs testing the effect of height-implied viewing geometry on detection hazard.

| Added term(s) | Key smooth term | edf | Ref.df | $\chi^2$ | p-value |
| --- | --- | --- | --- | --- | --- |
| Baseline distance | s(tend) | 8.815 | 10.44 | 715.6 | <0.001 |
| PAMM |  |  |  |  |  |
| Height | s(Height) | 2.068 | 2.204 | 6.078 | 0.057 |
| Height + tensor | s(Height) | 2.466 | 2.639 | 7.489 | 0.039 |
| interaction | ti(tend, Height) | 10.208 | 14.406 | 28.472 | 0.016 |

Pattern type significantly affected detection risk. Relative to the blotch pattern, the zigzag pattern was associated with a significantly lower detection hazard, indicating that zigzag-patterned models were more difficult to detect and therefore achieved better camouflage. By contrast, the banding, spot, and diamond patterns did not differ significantly from the blotch pattern, although the spot pattern showed a marginal trend towards reduced detection risk (Figure 2, Figure 3). Grey models had a significantly higher detection hazard than brown models, whereas sunny conditions significantly reduced detection hazard relative to cloudy conditions. Table 3 presents the full fixed-effect estimates for the distance- and angle-based baseline PAMMs.

**Figure 2.**
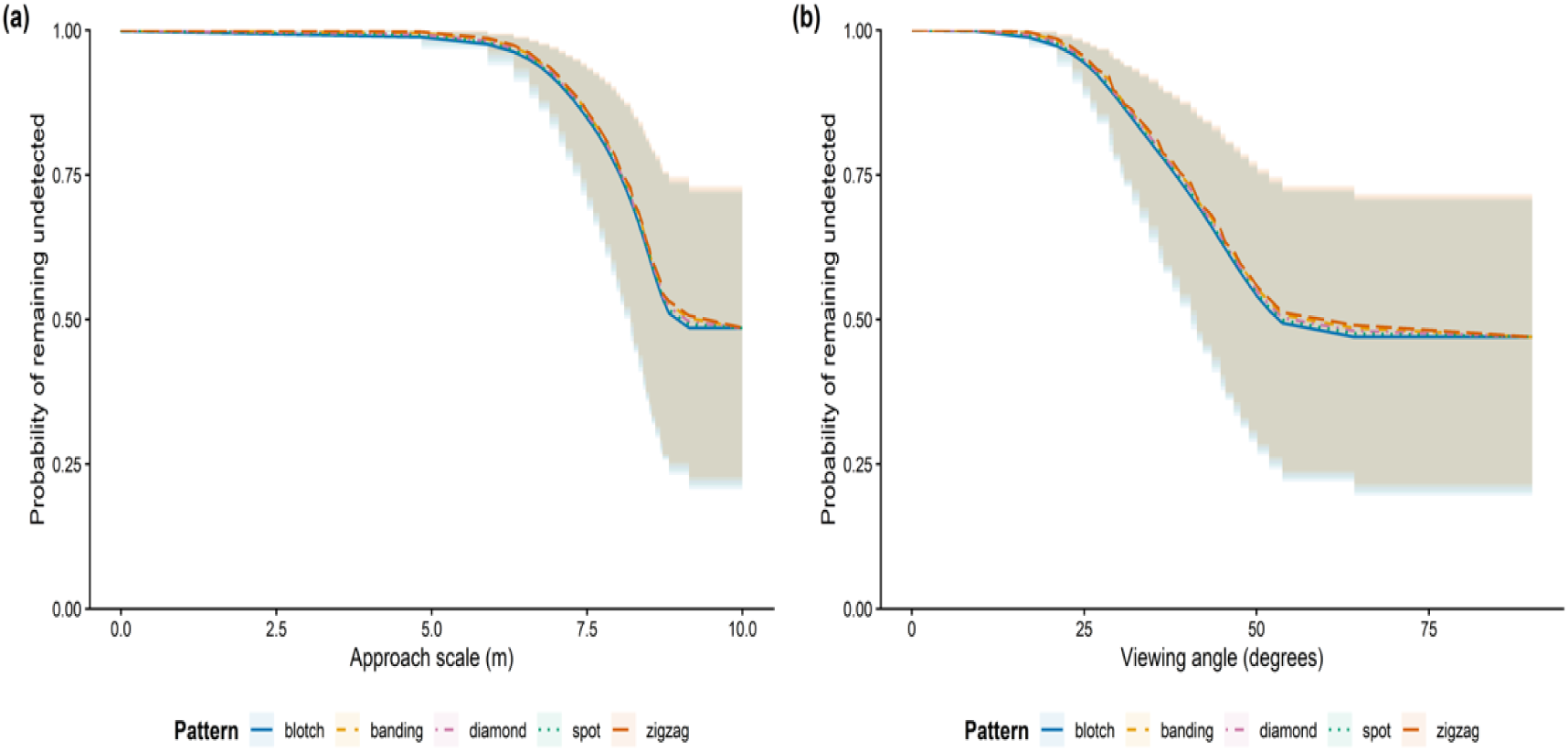
Predicted survival probabilities of snake models with different patterns from piece-wise exponential additive mixed models (PAMMs), predicted survival probabilities with 95% confidence intervals, based on N = 831 observations and 495 detection events. Lines represent predicted probabilities and shaded bands indicate the corresponding 95% confidence intervals.

**Figure 3.**
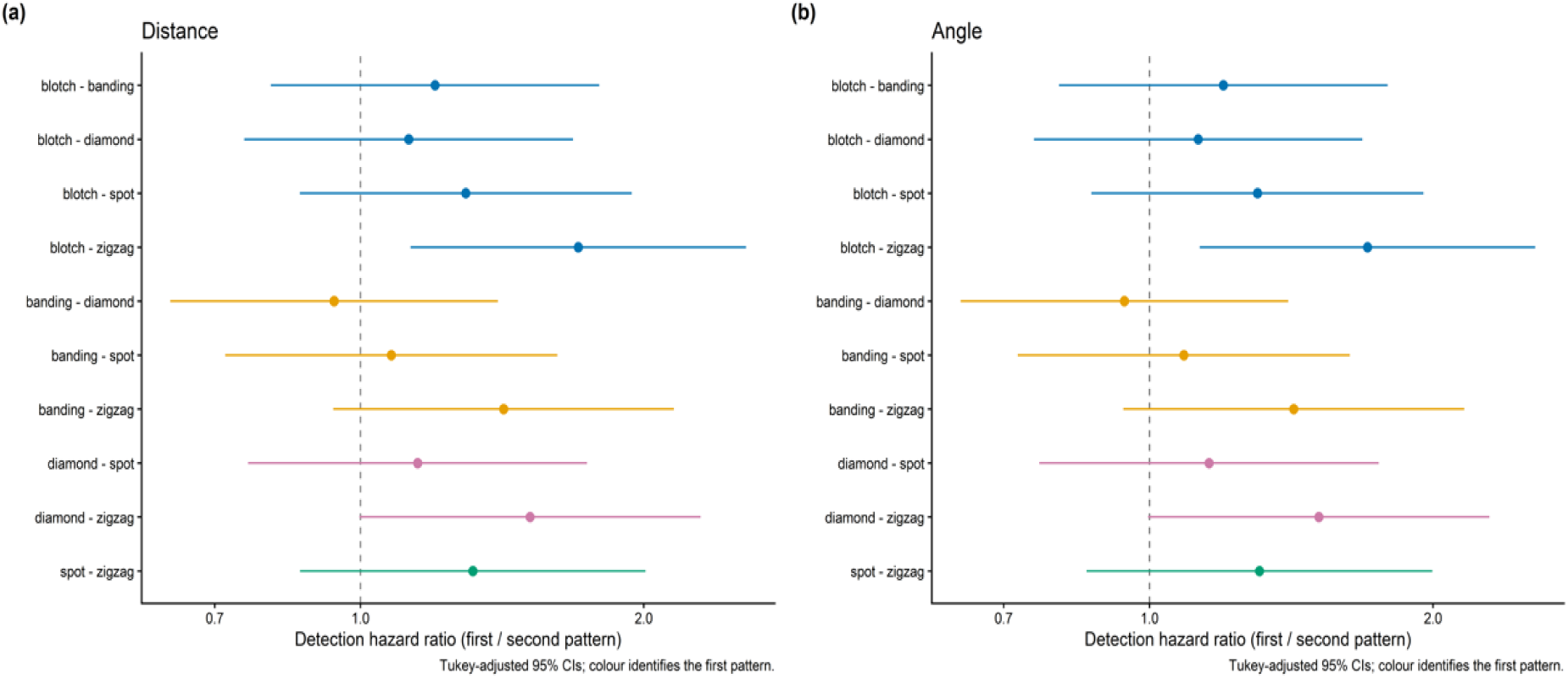
Pairwise comparisons of detection hazard among the five snake pattern treatments from (a) the distance-based and (b) the angle-based PAMMs. Points show hazard ratios and horizontal lines indicate Tukey-adjusted 95% confidence intervals for all ten pairwise comparisons within each model. Each hazard ratio compares the first pattern listed with the second; values below 1 indicate a lower detection hazard for the first pattern. The dashed vertical line indicates equal detection hazards (hazard ratio = 1); horizontal axes are logarithmic.

**Table 3.** Fixed effects from piece-wise exponential additive mixed models (PAMMs) describing detection hazard using distance and angle as alternative time scales.

|  | Distance |  |  |  | Angle |  |  |  |
| --- | --- | --- | --- | --- | --- | --- | --- | --- |
|  | Estimate | 95% CI | z | Pr(>\|z\|) | Estimate | 95% CI | z | Pr(>\|z\|) |
| Pattern: | -0.183 | [-0.471, | -1.238 | 0.216 | -0.180 | [-0.468, 0.108] | -1.225 | 0.221 |
| Banding |  | 0.105] |  |  |  |  |  |  |
| Pattern: | -0.258 | [-0.55, 0.034] | -1.732 | 0.083 | -0.264 | [-0.556, 0.028] | -1.775 | 0.076 |
| Spot |  |  |  |  |  |  |  |  |
| Pattern: | -0.118 | [-0.406, 0.17] | -0.800 | 0.424 | -0.119 | [-0.407, 0.169] | -0.808 | 0.419 |
| Diamond |  |  |  |  |  |  |  |  |
| Pattern: | -0.533 | [-0.829, - | -3.542 | <0.001 | -0.533 | [-0.827, -0.239] | -3.544 | <0.001 |
| Zigzag |  | 0.237] |  |  |  |  |  |  |
| Colour: | 0.477 | [0.263, 0.691] | 4.394 | <0.001 | 0.478 | [0.264, 0.692] | 4.399 | <0.001 |
| Grey |  |  |  |  |  |  |  |  |
| Weather: | -0.703 | [-1.007, - | -4.549 | <0.001 | -0.722 | [-1.018, -0.426] | -4.778 | <0.001 |
| Sunny |  | 0.399] |  |  |  |  |  |  |

Pairwise comparisons among pattern types showed that the zigzag pattern differed significantly from the blotch pattern, whereas the contrast between zigzag and diamond was marginally non-significant. No other pairwise differences were significant after adjustment for multiple comparisons (Table S3).

To test for pattern-specific hazard trajectories over distance, we fitted an extended model including pattern-specific smooth terms. However, this model did not improve fit relative to the simpler distance-based PAMM (ΔAIC = 6.842, Table S1), and none of the pattern-specific smooth terms were significant (Table S4). This suggests that patterns differed primarily in overall detection hazard rather than a strong difference in the shape of the hazard curve itself across distance.

### Robustness to angular scaling and the effect of observer height

Among detected models, viewing angles at detection ranged from 9.3° to 64.2°. The angle-based PAMM yielded the same main pattern effect as the distance-based model. In particular, the zigzag pattern again showed a significantly lower detection hazard than the blotch pattern, while banding, spot, and diamond did not differ significantly from blotch. Grey models remained more detectable than brown models, and sunny conditions again reduced detection hazard (Table 3). Pairwise contrasts were also similar to those obtained in the distance-based analysis, with a significant difference only between zigzag and blotch, and a marginal contrast between zigzag and diamond (Figure 2, Figure 3, Table S1). These results indicate that the lower detectability of the zigzag pattern was robust to whether the search process was represented using distance or angle.

However, unlike the distance-based analysis, the extended angle-based PAMM that included pattern-specific smooth terms showed a modest improvement in fit over the simpler model (Table S1). In this model, the pattern-specific smooths for blotch, spot, and diamond were significant, and the smooth for banding was marginally significant, whereas the smooth for zigzag was not significant (Table S4). This suggests that although the main ranking of patterns remained unchanged, expressing the search process on an angular scale revealed additional variation in how detection hazard changed across approach for some pattern types. Thus, the angle-based analysis supports the robustness of the main pattern result, while also indicating that viewing geometry may influence the detailed form of hazard trajectories.

To further investigate whether viewing geometry itself influenced detectability, we fitted supplementary distance-based PAMMs including observer height. A model including only a smooth main effect of height did not improve fit relative to the baseline distance model (AIC = 5225.726 vs 5223.020), and the smooth for height was only marginally significant (Table 2, Table S1). By contrast, when a tensor interaction between distance and height was included, model fit improved substantially (AIC = 5209.612), and both the main smooth for height and the interaction term were significant (Table 2, Table S1). These results indicate that observer height, and implied viewing geometry, did not simply shift detection risk by a constant amount, but modified how detection hazard changed during approach.

### Estimated camouflage effect based on image analysis

For the photographic analysis of snake models, we compared the effect of different patterns on edge disruption, chromatic contrast and achromatic contrast within the human visual system at distances of 5–15 m. The results of the LMM model showed that pattern, body colour, and their interaction all had significant effects on the GabRat value; chromatic contrast (ΔS) and luminance contrast (ΔS_Lum; Table S2).

Pairwise comparisons among different patterns revealed that the banding pattern exhibited the highest GabRat value compared to all other patterns. Additionally, the blotch pattern produced significantly lower GabRat values than the zigzag and diamond patterns, while the spot pattern exhibited significantly lower GabRat values than the diamond pattern and even less than the zigzag pattern (Table S3). Thus, under the standardised image-analysis conditions, pattern type strongly affected estimated edge disruption, with transverse or dorsal high-contrast markings generally producing higher GabRat values than more irregular or discrete lateral markings. Chromatic contrast differed statistically among patterns, with the diamond pattern showing the highest ΔS. However, the absolute differences among treatments were very small, and most pairwise differences excluding diamond pattern were all below 0.1 JND, indicating minimal practical differences in chromatic contrast (Table S3). This was expected because all pattern treatments used the same black markings and the same two base colours. By contrast, achromatic contrast showed clearer differences among patterns: spot and blotch models had the highest ΔS_Lum, diamond models had the lowest ΔS_Lum, and banding and zigzag models were intermediate and did not differ significantly from each other (Table S3). These results indicate that pattern design affected overall luminance contrast against the background, but the lower detectability of zigzag models cannot be explained by luminance contrast alone.

Comparisons of GabRat values at 2.5-meter distance intervals between 5 m and 15 m for the same pattern type showed that, to the human visual system model, adjacent distance intervals exhibited significant differences (Table S5). Furthermore, GabRat values increased with viewing distance (Figure 4), indicating stronger potential for pattern-outline interference at greater distances. Although the ΔS changed significantly between adjacent distance levels within the same pattern type, the relative changes are all below 1%, therefore probably too small to be distinguished by the human visual system, the significant deviation only exists on the statistical level. This indicates that as human observers approach the target, the measured edge disruption potential (GabRat) rapidly diminished, whereas the chromatic contrast remained largely unchanged as distance decreased.

**Figure 4.**
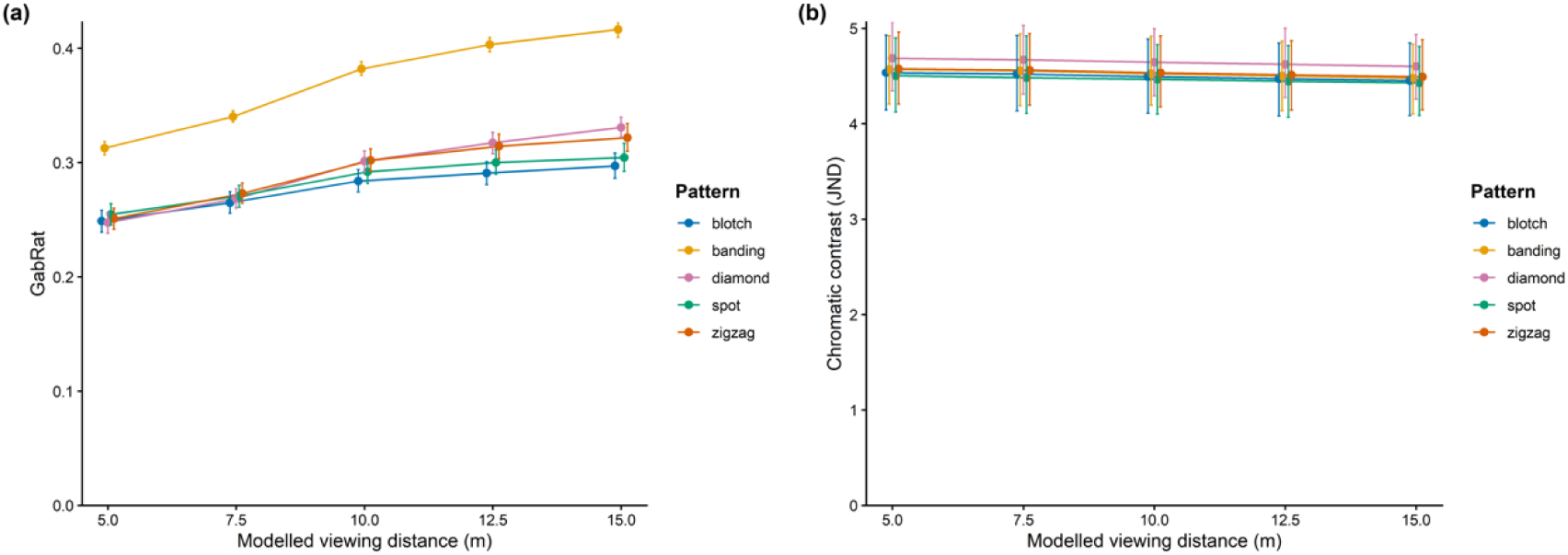
Image-based measures of (a) potential edge disruption (GabRat) and (b) chromatic contrast between snake models and the local background (ΔS, in just noticeable differences, JNDs) across modelled viewing distances for the human visual system. Points show means across placement sites and body colours, and error bars indicate bootstrap 95% confidence intervals. Lines connect means for each pattern.

## Discussion

The volunteer search results showed that zigzag patterning can enhance camouflage effectiveness, with zigzag-patterned models being detected at significantly shorter distances than blotch-patterned models and showing a marginal non-significant trend towards shorter detection distances than diamond-patterned models. Although the final overall detection rate did not differ among treatments, observers needed to be at a closer distance to detect snakes with zig-zags. Detection of the models did not increase gradually as volunteers got closer; instead, there was a steep rise in detection over part of the approach process. This suggests a threshold-like change in detectability. This pattern is consistent with a rapid reduction in camouflage effectiveness at closer viewing distances, it may also partly reflect observer search behaviour or changes in visibility caused by local vegetation. The present analyses support a cautious interpretation of viewing geometry. Re-expressing the search process in angular terms did not change the main conclusion that zigzag models had the lowest detection hazard, indicating that this pattern advantage is robust to geometric scale. At the same time, the angle-based model suggested that some pattern-specific hazard trajectories may differ across angular approach. Together, these findings suggest that detectability is influenced by viewing geometry, but do not support the stronger claim that angle is intrinsically more important than distance. The image analysis showed that all the patterns’ edge disruption effects increased with observer distance, while the contrast against the background remained stable with distance. This suggests that the distance-dependent camouflage effect change of these patterns is dominated by edge disruption rather than background matching. This is to be expected in our experiment since, beyond the difference in grey or brown base colour snake models, all snakes were the same colours.

The functional role—camouflage or conspicuous—of the snake patterns examined in this study is supported by evidence on both sides. Spot patterns including blotches and diamond patterns are considered to be cryptic in specific habitats which have background shadows and irregular objects (Allen et al., 2013; Jackson et al., 1976). However, in the racer (*Coluber constrictor*), individuals with spot patterns exhibit heightened aggression in response to predators, indicating a possible aposematic function (Creer, 2005). In our image analysis, the edge disruption of spot patterns declines markedly at close range. Such aggressive behaviour may offset this limitation, supporting Allen’s (2013) conclusion that pattern function is closely linked to behavioural traits. Banding patterns are commonly considered as aposematic (Brodie III, 1993; Campbell & Lamar, 1991; Savage & Slowinski, 1992), though some research suggests they may also serve a camouflage role (Pough, 1976). Our image analysis showed that the banding pattern perpendicular to body outline is highly contrasting and produces the highest GabRat values. Yet this strong potential for edge disruption did not yield corresponding camouflage benefits in the detection task. This may be because the regular repetition and high contrast bands that generate high GabRat values may also act as a cue for detection (Merilaita & Lind, 2006), and that high contrast can either confer camouflage or conspicuousness depending on the specific pattern arrangement. Moreover, such features are also characteristic of aposematic signals, and the balance between camouflage and warning functions may be further tuned by body coloration, which influences how the pattern interacts with the background. Thus, the same pattern properties can have opposing effects on detectability depending on context.

The shorter detection distances for zigzag models are consistent with recent image-based analyses suggesting that zigzag markings can generate strong predicted edge disruption under particular combinations of habitat, viewing distance and viewing angle, and the present study adds behavioural evidence that this visual effect can correspond to reduced detectability during active search (Li et al., 2026). Previous studies using predation simulation experiments with snake models have often interpreted the zigzag pattern primarily as aposematic, rather than as having a camouflage function (Niskanen & Mappes, 2005; J. Valkonen et al., 2011; Wüster et al., 2004). This appears to contradict the conclusions of the present study. However, by manipulating the backgrounds of their models, studies by Wüster et al. (2004) and Niskanen and Mappes (2005), excluded background matching as a potential camouflage mechanism for zigzag patterns, but did not exclude the possibility of edge disruption camouflage. It is also important to note that except for J. K. Valkonen et al. (2020) which used human volunteers, other studies (Niskanen & Mappes, 2005; J. Valkonen et al., 2011; Wüster et al., 2004) were based on measured bird attacks. Raptors view the snakes from above, and their perspective minimizes the influence of ground vegetation. While higher viewing angles intensify distance effects, they may also weaken edge disruption (Li et al., 2026), making aposematic signalling more important. Furthermore, these studies often compared the survival rates of patterns with potential camouflage or aposematic effects against those of stripe or uniform colour patterns. The aposematic signals may still benefit from being less conspicuous than other highly visible warning signals (J. A. Endler & Mappes, 2004). Therefore, rather than contradicting previous research, our findings may serve as a complement.

By manipulating pattern size (Bohlin et al., 2008), colour (Tullberg et al., 2005), spatial frequency (Barnett & Cuthill, 2014; Barnett et al., 2017), or directly evaluating the achromatic and chromatic contrast of targets against the background by image analysis and human detection (Barnett, Cuthill, et al., 2018; Barnett, Michalis, et al., 2018; Caro, Stankowich, Kiffner, & Hunter, 2013), distance-dependent defensive coloration research usually focuses on background matching. But beyond colour matching, false edges generated by putative disruptive patterns are also affected by the distance and light conditions according to our results. Due to limits in observer visual acuity, pattern elements may blend into an average colour when observed from distance, making discrimination harder when contrast is low (Marshall, 2000). Given the spatial scale of the pattern elements and of human visual acuity used in the experiment, complete distance-dependent pattern blending due to limited visual acuity was unlikely within the range tested. However, high-contrast patterns may emphasize the pattern to be more conspicuous than the outline, allowing disruptive elements to be resolved before the body boundary. Under the direct light condition, darker patterns may blend with shadows and enhance the false edges, thereby contributing distance-dependent defensive coloration through edge disruption.

Supplementary analyses further indicated that observer height can modify detection hazard over distance. This effect likely arises because environmental obstructions and barriers (e.g., vegetation) partially conceal body contours or recognizable features, which are key cues for target identification, thereby enhancing their camouflage (Hancock et al., 2023; Stevens et al., 2017; Tvardíková & Fuchs, 2010). Due to the presence of these obstructions and barriers, individuals at different viewing heights experience different levels of visual obstruction. At the same distance, observers with a higher viewing angle are better able to discern a target (Hancock et al., 2023). Raptors, often the primary predators of snakes, could face reduced hunting success if snake patterns exhibit distance-dependent camouflage effects, which may help mitigate predation pressure from aerial predators.

Beyond the external environment, the inherent geometry of pattern distribution on the animal’s own body surface introduces a second layer of viewing-angle dependency. Symmetrical patterns can serve as cues for predators, compromising camouflage effectiveness, and have a stronger impact on background-matching camouflage than on edge disruption (Cuthill, Hiby, & Lloyd, 2006; Cuthill, Stevens, Windsor, & Walker, 2006), whereas asymmetrical and variable patterns are more effective in avoiding detection (Merilaita & Lind, 2006). The same target may appear differently depending on body pattern distribution, leading to variations in conspicuousness and perceptibility for observers at different positions. For instance, the detectability of hoverfly mimicry (Baker et al., 2024) and the conspicuousness of poison frogs (McEwen, Yeager, Kinley, Anderson, & Barnett, 2024) shift with viewing angle. Such detectability differences may lead to multiple functions for different signal receivers. Moreover, animals may also adjust their posture to alter their appearance, minimizing unfavourable pattern features and improving camouflage (Barbosa, Allen, Mäthger, & Hanlon, 2012; C. Kang, Stevens, Moon, Lee, & Jablonski, 2015; C. K. Kang, Moon, Lee, & Jablonski, 2012). For snakes, these viewing-angle effects may also be relevant to encounters with natural terrestrial predators, which can view the same body pattern from different elevations and approach directions, potentially altering the visibility of pattern elements and body contours.

The threshold-like increase in detection during observer approach may also have been influenced by observers preferentially searching within a particular distance range ahead of them. Partial occlusion by local vegetation was not quantified but may have contributed to abrupt changes in visibility. Variation in human observer height also reflects a relatively restricted range of terrestrial viewing geometries and should not be taken to represent the full range of viewpoints experienced by terrestrial and aerial predators of snakes. Although we standardised the approximate size and longitudinal spacing of repeated markings as far as possible, the pattern treatments were not matched for total pattern area or black coverage. The treatments should therefore be interpreted as whole-pattern phenotypes, in which marking shape, arrangement, and coverage varied together. However, the brightness contrast analysis suggested that luminance contrast alone is unlikely to explain the lower detectability of zigzag models.

Future work could manipulate luminance contrast, pattern coverage and pattern arrangement independently to separate their effects on edge disruption and detectability more directly. Complementary studies could also integrate controlled obstacles and viewing angles into camouflage studies, while considering observer search behaviour, especially for non-uniformly distributed patterns on 3D objects. Virtual modelling (Miller, Hogan, & Stoddard, 2022) and multi-angle scanning with calibrated cameras may refine investigations in this area. Overall, these findings indicate a distance-dependent camouflage effect shaped by angular scaling and observer height. However, rather than demonstrating that viewing angle outweighs distance, the analyses suggest that distance, angular scaling, and observer height jointly influence how pattern detectability changes during approach.

## Data Availability

The anonymized data underlying the statistical analyses, the R code used for data processing, statistical analyses and figure generation, and a comprehensive README file are provided with this submission. Image processing was conducted using the Multispectral Image Calibration and Analysis Toolbox in ImageJ, no code was generated for that stage.

## Declaration of Generative AI and AI-assisted Technologies in the Manuscript Preparation Process

During the preparation of this work, the authors used ChatGPT (OpenAI) to assist with language editing and code debug. After using this tool, the authors reviewed and edited the content as needed and take full responsibility for the content of the article.

## Author contributions

**Zixiang Li:** Conceptualization; Methodology; Investigation; Data curation; Formal analysis; Software; Visualization; Funding acquisition; Writing – original draft; Writing – review & editing.

**Laura A. Kelley:** Methodology; Supervision; Writing – review & editing.

**Martin Stevens:** Conceptualization; Methodology; Supervision; Project administration; Writing – review & editing.

