## Supplementary material for "Viewing geometry and pattern type shape the detectability of snakes": Table S1

| **Model comparison of PAMMs used in field detection analyses** | | | | | |
| --- | --- | --- | --- | --- | --- |
| Analysis set | Model | Description | df | AIC | ΔAIC |
| Distance-based models | m0 | baseline distance-based PAMM | 58.535 | 5223.020 |  |
|  | m1 | pattern-specific smooth | 73.317 | 5229.862 | 6.842 |
| Angle-based models | m0_angle | baseline angle-based PAMM | 56.143 | 5413.179 |  |
|  | m1_angle | pattern-specific smooth | 61.571 | 5407.604 | -5.575 |
| Height-based supplementary models | m0 | baseline distance-based PAMM | 58.535 | 5223.020 |  |
|  | m_h_dist | includes a smooth main effect of Height | 58.352 | 5225.726 | 2.706 |
|  | m_h_ti | includes a tensor interaction ti(tend, Height) | 71.717 | 5209.612 | -13.408 |

Table S2.

| **Effect of pattern and colour on the camouflage in LMM model based on snake replica photograph** | | | | | | | | |
| --- | --- | --- | --- | --- | --- | --- | --- | --- |
|  | Fixed effect | Sum Sq | Mean Sq | NumDF | DenDF | F value | Pr(>F) |  |
| GabRat | Pattern | 1.752 | 0.438 | 4 | 1461 | 352.256 | <0.001 | *** |
|  | Colour | 0.830 | 0.830 | 1 | 1461 | 667.692 | <0.001 | *** |
|  | Pattern:Colour | 0.087 | 0.022 | 4 | 1461 | 17.463 | <0.001 | *** |
| dS | Pattern | 5.61 | 1.40 | 4 | 1461 | 44.229 | <0.001 | *** |
|  | Colour | 1100.92 | 1100.92 | 1 | 1461 | 34728.121 | <0.001 | *** |
|  | Pattern:Colour | 7.94 | 1.99 | 4 | 1461 | 62.637 | <0.001 | *** |
| dS_Lum | Pattern | 135.09 | 33.77 | 4 | 1461 | 311.96 | <0.001 | *** |
|  | Colour | 1776.49 | 1776.49 | 1 | 1461 | 16410.00 | <0.001 | *** |
|  | Pattern:Colour | 59.61 | 14.90 | 4 | 1461 | 137.65 | <0.001 | *** |

Table S3a.

| **Pairwise comparisons among pattern treatments in the field-detection PAMMs.** | | | | | | | | |
| --- | --- | --- | --- | --- | --- | --- | --- | --- |
| Contrast | Distance | | | | Angle | | | |
|  | estimate | 95% CI | t.ratio | p.value | estimate | 95% CI | t.ratio | p.value |
| blotch-banding | 0.183 | [-0.105, 0.471] | 1.238 | 0.729 | 0.180 | [-0.108, 0.468] | 1.225 | 0.737 |
| blotch-spot | 0.258 | [-0.034, 0.55] | 1.732 | 0.414 | 0.264 | [-0.028, 0.556] | 1.775 | 0.388 |
| blotch-diamond | 0.118 | [-0.17, 0.406] | 0.800 | 0.931 | 0.119 | [-0.169, 0.407] | 0.808 | 0.928 |
| blotch-zigzag | 0.533 | [0.239, 0.827] | 3.542 | **0.004** | 0.533 | [0.239, 0.827] | 3.544 | **0.004** |
| banding-spot | 0.076 | [-0.216, 0.368] | 0.507 | 0.987 | 0.084 | [-0.208, 0.376] | 0.563 | 0.980 |
| banding-diamond | -0.065 | [-0.353, 0.223] | -0.439 | 0.992 | -0.062 | [-0.350, 0.226] | -0.419 | 0.994 |
| banding-zigzag | 0.350 | [0.050, 0.650] | 2.293 | 0.147 | 0.353 | [0.053, 0.653] | 2.308 | 0.142 |
| spot-diamond | -0.140 | [-0.438, 0.158] | -0.921 | 0.889 | -0.145 | [-0.443, 0.153] | -0.955 | 0.875 |
| spot-zigzag | 0.275 | [-0.029, 0.579] | 1.774 | 0.389 | 0.269 | [-0.035, 0.573] | 1.734 | 0.413 |
| diamond-zigzag | 0.415 | [0.115, 0.715] | 2.711 | 0.052 | 0.414 | [0.114, 0.714] | 2.707 | 0.053 |

**Table S3b.**

|  | **Pairwise comparisons among pattern treatments in the image-analysis models.** | | | | | | | | | | | |
| --- | --- | --- | --- | --- | --- | --- | --- | --- | --- | --- | --- | --- |
| Contrast | GabRat | | | | dS | | | | dS_Lum | | | |
|  | estimate | 95% CI | t.ratio | p.value | estimate | 95% CI | t.ratio | p.value | estimate | 95% CI | t.ratio | p.value |
| blotch-banding | -0.094 | [-0.1, -0.088] | -32.58 | **<0.001** | -0.032 | [-0.061, -0.003] | -2.175 | 0.2977 | 0.498 | [0.445, 0.551] | 18.534 | <0.001 |
| blotch-spot | -0.007 | [-0.013, -0.001] | -2.519 | 0.119 | 0.03 | [0.001, 0.059] | 2.079 | 0.378 | -0.054 | [-0.107, -0.001] | -1.999 | 0.457 |
| blotch-diamond | -0.016 | [-0.022, -0.01] | -5.527 | **<0.001** | -0.15 | [-0.179, -0.121] | -10.321 | **<0.001** | 0.723 | [0.67, 0.776] | 26.902 | <0.001 |
| blotch-zigzag | -0.015 | [-0.021, -0.009] | -5.313 | **<0.001** | -0.04 | [-0.069, -0.011] | -2.754 | 0.0597 | 0.440 | [0.387, 0.493] | 16.382 | <0.001 |
| banding-spot | 0.087 | [0.081, 0.093] | 30.061 | **<0.001** | 0.062 | [0.033, 0.091] | 4.254 | **<0.001** | -0.552 | [-0.605, -0.499] | -20.533 | <0.001 |
| banding-diamond | 0.078 | [0.072, 0.084] | 27.053 | **<0.001** | -0.118 | [-0.147, -0.089] | -8.145 | **<0.001** | 0.225 | [0.172, 0.278] | 8.368 | <0.001 |
| banding-zigzag | 0.079 | [0.073, 0.085] | 27.267 | **<0.001** | -0.008 | [-0.037, 0.021] | -0.578 | 1 | -0.058 | [-0.111, -0.005] | -2.152 | 0.3156 |
| spot-diamond | -0.009 | [-0.015, -0.003] | -3.008 | **0.027** | -0.18 | [-0.209, -0.151] | -12.4 | **<0.001** | 0.776 | [0.723, 0.829] | 28.901 | <0.001 |
| spot-zigzag | -0.008 | [-0.014, -0.002] | -2.794 | 0.053 | -0.07 | [-0.099, -0.041] | -4.833 | **<0.001** | 0.494 | [0.441, 0.547] | 18.381 | <0.001 |
| diamond-zigzag | 0.001 | [-0.005, 0.007] | 0.214 | 1 | 0.11 | [0.081, 0.139] | 7.567 | **<0.001** | -0.283 | [-0.336, -0.23] | -10.520 | <0.001 |

Table S4.

| **Approximate significance of pattern-specific smooth terms in extended distance- and angle-based PAMMs** | | | | | |
| --- | --- | --- | --- | --- | --- |
| Analysis set | Smooth term | edf | Ref.df | χ² | p-value |
| Distance-based extended PAMM | s(tend) | 8.853 | 10.470 | 358.018 | <0.001 |
|  | s(tend):Patternblotch | 1.000 | 1.000 | 1.421 | 0.233 |
|  | s(tend):Patternbanding | 1.425 | 1.733 | 0.497 | 0.749 |
|  | s(tend):Patternspot | 0.000 | 0.000 | 0.000 | 0.995 |
|  | s(tend):Patterndiamond | 2.128 | 2.739 | 2.713 | 0.477 |
|  | s(tend):Patternzigzag | 2.987 | 3.809 | 6.552 | 0.137 |
|  | s(Volunteer) | 20.830 | 27.000 | 98.664 | <0.001 |
|  | s(Point) | 20.090 | 29.000 | 68.390 | <0.001 |
| Angle-based extended PAMM | s(tend) | 6.850 | 8.473 | 210.723 | <0.001 |
|  | s(tend):Patternblotch | 1.001 | 1.001 | 11.279 | 0.001 |
|  | s(tend):Patternbanding | 1.002 | 1.004 | 3.439 | 0.064 |
|  | s(tend):Patternspot | 1.000 | 1.000 | 6.091 | 0.014 |
|  | s(tend):Patterndiamond | 1.483 | 1.841 | 13.693 | 0.003 |
|  | s(tend):Patternzigzag | 0.000 | 0.000 | 0.000 | 0.500 |
|  | s(Volunteer) | 20.280 | 27.000 | 87.141 | <0.001 |
|  | s(Point) | 20.230 | 29.000 | 68.944 | <0.001 |

**Table S5.**

| **Comparison of camouflage effect of each pattern between different distance levels** | | | | | | | | | | | | | |
| --- | --- | --- | --- | --- | --- | --- | --- | --- | --- | --- | --- | --- | --- |
|  |  | dS | | | | | | GabRat | | | | | |
| Pattern | Comparison | Mean_Diff | CI_Lower | CI_Upper | t-value | p-value | Relative Change (%) | Mean_Diff | CI_Lower | CI_Upper | t-value | p-value | Relative Change (%) |
| blotch | 5 vs 7.5 | 0.015 | 0.008 | 0.021 | 4.322 | <0.001 | 0.322 | -0.016 | -0.019 | -0.013 | -12.614 | <0.001 | -6.216 |
|  | 7.5 vs 10 | 0.028 | 0.019 | 0.036 | 6.180 | <0.001 | 0.611 | -0.019 | -0.022 | -0.016 | -12.617 | <0.001 | -6.911 |
|  | 10 vs 12.5 | 0.022 | 0.016 | 0.028 | 7.509 | <0.001 | 0.500 | -0.007 | -0.009 | -0.005 | -7.581 | <0.001 | -2.457 |
|  | 12.5 vs 15 | 0.019 | 0.014 | 0.025 | 7.554 | <0.001 | 0.435 | -0.006 | -0.008 | -0.004 | -6.466 | <0.001 | -2.116 |
| banding | 5 vs 7.5 | 0.011 | 0.003 | 0.019 | 2.757 | 0.008 | 0.235 | -0.028 | -0.030 | -0.025 | -22.595 | <0.001 | -8.473 |
|  | 7.5 vs 10 | 0.032 | 0.024 | 0.040 | 8.105 | <0.001 | 0.713 | -0.042 | -0.045 | -0.039 | -30.235 | <0.001 | -11.564 |
|  | 10 vs 12.5 | 0.024 | 0.019 | 0.030 | 9.025 | <0.001 | 0.542 | -0.021 | -0.023 | -0.019 | -23.511 | <0.001 | -5.404 |
|  | 12.5 vs 15 | 0.021 | 0.017 | 0.026 | 9.621 | <0.001 | 0.470 | -0.013 | -0.015 | -0.011 | -14.271 | <0.001 | -3.227 |
| spot | 5 vs 7.5 | 0.021 | 0.014 | 0.028 | 6.322 | <0.001 | 0.471 | -0.016 | -0.019 | -0.014 | -12.809 | <0.001 | -6.202 |
|  | 7.5 vs 10 | 0.016 | 0.009 | 0.023 | 4.422 | <0.001 | 0.351 | -0.021 | -0.024 | -0.018 | -12.538 | <0.001 | -7.474 |
|  | 10 vs 12.5 | 0.020 | 0.014 | 0.026 | 6.779 | <0.001 | 0.454 | -0.008 | -0.010 | -0.006 | -8.065 | <0.001 | -2.779 |
|  | 12.5 vs 15 | 0.018 | 0.013 | 0.023 | 7.417 | <0.001 | 0.400 | -0.004 | -0.007 | -0.002 | -3.636 | 0.001 | -1.411 |
| diamond | 5 vs 7.5 | 0.015 | 0.008 | 0.021 | 4.654 | <0.001 | 0.319 | -0.021 | -0.023 | -0.019 | -17.577 | <0.001 | -8.147 |
|  | 7.5 vs 10 | 0.026 | 0.018 | 0.034 | 6.634 | <0.001 | 0.560 | -0.033 | -0.035 | -0.030 | -24.219 | <0.001 | -11.501 |
|  | 10 vs 12.5 | 0.022 | 0.017 | 0.028 | 7.696 | <0.001 | 0.484 | -0.016 | -0.018 | -0.014 | -17.193 | <0.001 | -5.231 |
|  | 12.5 vs 15 | 0.020 | 0.015 | 0.026 | 7.601 | <0.001 | 0.441 | -0.013 | -0.015 | -0.011 | -12.990 | <0.001 | -4.119 |
| zigzag | 5 vs 7.5 | 0.013 | 0.006 | 0.020 | 3.866 | <0.001 | 0.289 | -0.022 | -0.025 | -0.019 | -15.230 | <0.001 | -8.434 |
|  | 7.5 vs 10 | 0.028 | 0.020 | 0.037 | 6.593 | <0.001 | 0.617 | -0.029 | -0.032 | -0.026 | -18.209 | <0.001 | -10.059 |
|  | 10 vs 12.5 | 0.022 | 0.016 | 0.028 | 7.219 | <0.001 | 0.491 | -0.012 | -0.015 | -0.010 | -10.171 | <0.001 | -4.044 |
|  | 12.5 vs 15 | 0.020 | 0.014 | 0.025 | 7.573 | <0.001 | 0.436 | -0.007 | -0.010 | -0.005 | -5.410 | <0.001 | -2.332 |
